# Nested species distribution models of *Chlamydiales* in tick host *Ixodes ricinus* in Switzerland

**DOI:** 10.1101/2020.05.26.118216

**Authors:** Estelle Rochat, Séverine Vuilleumier, Sebastien Aeby, Gilbert Greub, Stéphane Joost

## Abstract

The tick *Ixodes ricinus* is the vector of various pathogens, including *Chlamydiales* bacteria, potentially causing respiratory infections. In this study, we modelled the spatial distribution of *I. ricinus* and associated *Chlamydiales* over Switzerland from 2009 to 2019. We used a total of 2293 ticks and 186 *Chlamydiales* occurrences provided by a Swiss Army field campaign, a collaborative smartphone application and a prospective campaign. For each tick location, we retrieved from Swiss federal datasets the environmental factors reflecting the topography, climate and land cover. We then used the Maxent modelling technique to estimate the suitability for *I. ricinus* and to subsequently build the nested niche of *Chlamydiales* bacteria. Results indicate that *I. ricinus* high habitat suitability is determined by higher temperature and vegetation index (NDVI) values, lower temperature during driest months and a higher percentage of artificial and forests areas. The performance of the model was increased when extracting the environmental variables for a 100 m-radius buffer around the sampling points and when considering the data over the two years previous sampling date. For *Chlamydiales* bacteria, the suitability was favoured by lower percentage of artificial surfaces, driest conditions, high precipitation during coldest months and short distances to wetlands. From 2009 to 2018, we observed an extension of tick and *Chlamydiales* suitable areas, associated with a shift towards higher altitude. The importance to consider spatio-temporal variations of the environmental conditions for obtaining better prediction was also demonstrated.

**Importance:** *Ixodes ricinus* is the vector of pathogens, including the agent of Lyme disease, the tick borne encephalitis virus and the less known *Chlamydiales* bacteria at the origin of some respiratory infections. In this study, we identified the environmental factors influencing the presence of *I. ricinus* and *Chlamydiales* in Switzerland and generated maps of their distribution from 2009 to 2018. We found an important expansion of suitable areas for both the tick and the bacteria during the last decade. Results provided also the environmental factors that determine the presence of *Chlamydiales* within ticks. Distribution maps as generated here are expected to bring valuable informations for decision-makers to control tick-borne diseases in Switzerland and establish prevention campaigns. The methodological framework presented could be used to predict the distribution and spread of other host-pathogen couples, to identify environmental factors driving their distribution and to develop control or prevention strategies accordingly.

## Introduction

*Ixodes ricinus* is the most common tick species in Switzerland and is known to be the vector of many pathogens, including the tick-borne encephalitis virus and the bacteria *Borrelia burgdoferi*, agent of the Lyme disease (Mermod *et al.*, 1973; Aeschlimann *et al.*, 1986). In 2015, Pilloux *et al.* showed that *I. ricinus* may also have a role of vector and even reservoir for *Chlamydiales* bacteria, especially *Rhabdochlamydiaceae* and *Parachlamydiaceae*. *Chlamydiales* is an order of strict intracellular bacteria containing various bacterial pathogens or emerging pathogens associated with serious diseases for humans and animals, including respiratory tract infections and miscarriage (Corsaro and Greub, 2006; Greub, 2009; Borel *et al.*, 2018). *Parachlamydiaceae* have been largely associated to free-living amoebae (Corsaro *et al.*, 2009, 2010) and are considered as emerging agents of pneumonia in humans (Lamoth and Greub, 2010a, 2010b). They have also been associated with miscarriage in ruminants (Borel *et al.*, 2007; Deuchande *et al.*, 2010) and have been documented in roe deer and red deer, as well as in some rodents (Regenscheit *et al.*, 2012; Stephan *et al.*, 2014). *Rhabdochlamydiaceae* have been mainly described associated to arthropods, including *Porcellio scaber, Blatta orientalis* and *Ixodes ricinus* (Kostanjsek *et al.*, 2004; Corsaro *et al.*, 2007; Pillonel *et al.*, 2019). The pathogenic role of *Rhabdochlamydiaceae* is still largely unknown, but suspected to cause newborn infections (Lamoth *et al.*, 2009) and respiratory complications such as pneumonia (Lamoth *et al.*, 2011).

Considering the potential threat to human health caused by pathogens associated with the tick *Ixodes ricinus*, studies already investigated the influence of environmental factors on its presence or density. They showed that the distribution and activity of *I. ricinus* is mainly influenced by temperature and humidity (Aeschlimann, 1972; Perret *et al.*, 2000, 2003; McCoy and Boulanger, 2015). Indeed, this tick species is prone to desiccation and a relative humidity between 70 to 80% close to the soil is necessary for its survival (Aeschlimann, 1972; Kahl and Alidousti, 1997; Perret *et al.*, 2000). Its most favourable habitats may therefore be vegetation types able to maintain a high humidity level close to the soil such as woodlands with thick vegetation litter (Aeschlimann, 1972; Lindgren *et al.*, 2006; McCoy and Boulanger, 2015).

In Switzerland, several studies analysed the impact of environmental conditions on the activity or density of *Ixodes ricinus*. An early study done by Aeschlimann *et al.* (1972) indicated that *I. ricinus* distribution is mainly limited by the presence of a favourable vegetation cover, with a relative humidity close or superior to 80% and an altitude inferior to 1500 m. Perret *et al.* (2000) showed that the questing activity of ticks takes place from a temperature of 7°C and Hauser *et al.* (2018) indicated that questing activity is largely reduced when the temperature exceeds 27°C. Jouda *et al.* (2004) showed that in the region of Neuchâtel, the density of ticks decrease with altitude, which was confirmed by Gern *et al.* (2008). However, this relationship was found opposite in the Alps (Valais), which they explained by drier conditions at lower altitude.

Bacteria communities within ticks are also known to be influenced by environmental conditions, notably through a modification of the tick density, the tick behaviour or the vector-host interactions (Carpi *et al.*, 2011; Ehrmann *et al.*, 2018; Aivelo *et al.*, 2019). For example, *B. burgdorferi* is most likely found at lower altitude (Gern *et al.*, 2008), infect more ticks collected in forests than in pastures (Halos *et al.*, 2010; Ehrmann *et al.*, 2018), and may be favoured by the forest fragmentation (Halos *et al.*, 2010; Roome *et al.*, 2018) while *Rickettsia* bacteria may be more prevalent in ticks in pasture sites showing a shrubby vegetation and a medium forest fragmentation (Halos *et al.*, 2010). Environmental factors might provide us with critical information for bacteria distribution and thus potential threats to human. However, nothing has been investigated regarding *Chlamydiales* bacteria yet.

Most studies described above analysed the impact of environmental factors on the density or questing activity of ticks. None modelled across years the spatial distribution of *Ixodes ricinus* habitat suitability at the Swiss scale nor the distribution of the *Chlamydiales* bacteria. In our study, we therefore aimed to build a model estimating the spatial distribution of the *I. ricinus* species from 2009 to 2019 in all Switzerland using the Maxent modelling technique. Beside, we also investigated, for the first time, the ecological factors that determine the distribution of *Chlamydiales* bacteria and the environmental factors that influence the presence of this bacteria within its tick host.

Modelling of *I. ricinus* distribution with Maxent has already been done at the scale of Europe (Porretta *et al.*, 2013), for an area including Europe, North Africa and Middle East (Alkishe *et al.*, 2017) and in Romania (Domsa *et al.*, 2018). Environmental data used in these studies were extracted from Worldclim climatic data at a spatial resolution of 30 arc-second (approximately 1 km). These data summarized climatic conditions from 1950 to 2000. Therefore in these studies as in many others (Williams *et al.*, 2015; Raghavan *et al.*, 2016, 2019, 2020; Sage *et al.*, 2017; Minigan *et al.*, 2018; Soucy *et al.*, 2018; Eisen *et al.*, 2018; Hadgu *et al.*, 2019) environmental data were extracted at a resolution that did not match the species ecology and more importantly the environmental conditions at sampling dates. Our goals were thus first to build a model of higher spatial resolution (100 m) for Switzerland and second to use recent climatic data to characterize in detail the distribution of *Ixodes ricinus* and its associated *Chlamydiales* bacterial pathogen over Switzerland from 2009 to 2019. To better understand the importance of the environmental conditions surrounding the sampling points, and the conditions preceding sampling date, we analysed the performance of the model 1) across buffer zones around the sampling point and 2) through different period of time before the sampling date. Finally, we investigated the potential to use the Maxent modelling to estimate the nested niche of a parasite within the ecological niche of its host.

## Material and Methods

Species distribution can be modelled with various methods that use either records of presence and absence of the species or only presences (Elith *et al.*, 2006; Tsoar *et al.*, 2007; Huerta and Peterson, 2008; Norberg *et al.*, 2019). Among them, the Maxent algorithm (Phillips *et al.*, 2006) using presence records only has been shown to perform particularly well as compared to other presence-only modelling methods (Elith *et al.*, 2006; Huerta and Peterson, 2008). We thus chose to use this model to determine the potential ecological niche of *Ixodes ricinus* and its associated *Chlamydiales* bacterial pathogen over Switzerland. The various steps of the method detailed in the paragraphs below are summarised on a Figure in Suppl. File 2.

### Ticks and bacteria occurrences data

Data regarding tick occurrences were obtained from three different sources. First, ticks were collected by a field campaign conducted by the **Swiss Army** from 21^st^ of April to 13^th^ of July 2009. During this campaign, 172 forests were sampled with convenience sampling in forests in altitude lower than 1,500 m. 62,889 ticks were collected by flagging low vegetation using a white-cloth. The ticks were then aggregated into 8’534 pools of 5 to 10 ticks (5 nymphs or 10 adults) and each pool was analysed for the presence of *Chlamydiales* DNA by using a pan-*Chlamydiales* real-time qPCR as described by Pilloux *et al.* (2015), after extracting the DNA as described by Gäumann *et al.* (2010). Among the 8,534 pools, 543 were positive (6.4%) and they were located in 118 out of the 172 sampling sites (68.6%).

Second, data were obtained from the collaborative smartphone application “**Tick Prevention**” (zecke-tique-tick.ch) developed by A&K Strategy GmbH, a Spin-off from the Zurich University of Applied Sciences (ZHAW) in which users can indicate tick locations on a map. The application was launched in February 2015 and by the end of December 2019, 29 153 locations of tick’s observations were available in Switzerland. To each observation a spatial accuracy is assigned depending on the scale (zoomed area) to which the observation was reported by the user. For our analysis, only observations with a spatial accuracy equal or higher to 100 m and only data collected from March to October were used. The final dataset corresponded to 5 781 tick’s locations. Moreover, since January 2017, users bitten by a tick can send the tick removed from their body to the national centre for tick-transmitted diseases (NRZK, www.labor-spiez.ch). The ticks received are analysed by three different laboratories for detecting the presence of various bacteria, including *Chlamydiales*. In April 2019, 554 ticks from 506 sites were received and sequenced, among which 21 ticks (3.79%) were positive for *Chlamydiales* bacteria and were located in 19 sites (3.75%).

Finally, to increase the number of data, especially regarding *Chlamydiales* occurrences, a **prospective campaign** was conducted by the authors from 11^th^ of May to 24^th^ of June 2018. During this campaign, 95 sites were visited, mainly in west Switzerland. Those sites were chosen in areas predicted to be favourable for the presence of ticks based on a pre-analysis of the two other datasets, and such to maximise the environmental variability between visited sites (see Suppl. File 1 for more details). Whenever possible, three ticks were collected in each site, by dragging a white-cloth over the soil. For some sites however, only one or two ticks could be found. Eventually, the campaign allowed the collection of 256 ticks, each of which were placed in a sterile tube and kept at 4°C before being sent to the laboratory to be analysed for the presence of *Chlamydiales* bacteria. In the laboratory, the ticks were washed once with 70% ethanol and twice with PBS. DNA was extracted using the NucleoSpin DNA Insect Kit (Macherey-Nagel) with NucleoSpin Bead Tubes Type E and MN Bead Tube Holder in combination with the Vortex-Genie 2. Manufacturer’s protocol was slightly adapted by performing disruption during 20 min followed by a 2h incubation at 56°C in order to allow proteinase K digestion. DNA was then analysed using the pan-*Chlamydiales* qPCR developed by Lienard *et al.* (2011). A tick was considered as positive for the presence of *Chlamydiales* if either the two replicates were positive or if one of the two was highly positive (CT value < 35). As a result, 72 out of the 256 ticks were positive (28.13%), in 51 out of 95 sites (53.6%).

The characteristics of each dataset are summarized in Table 1.

**Table 1:**
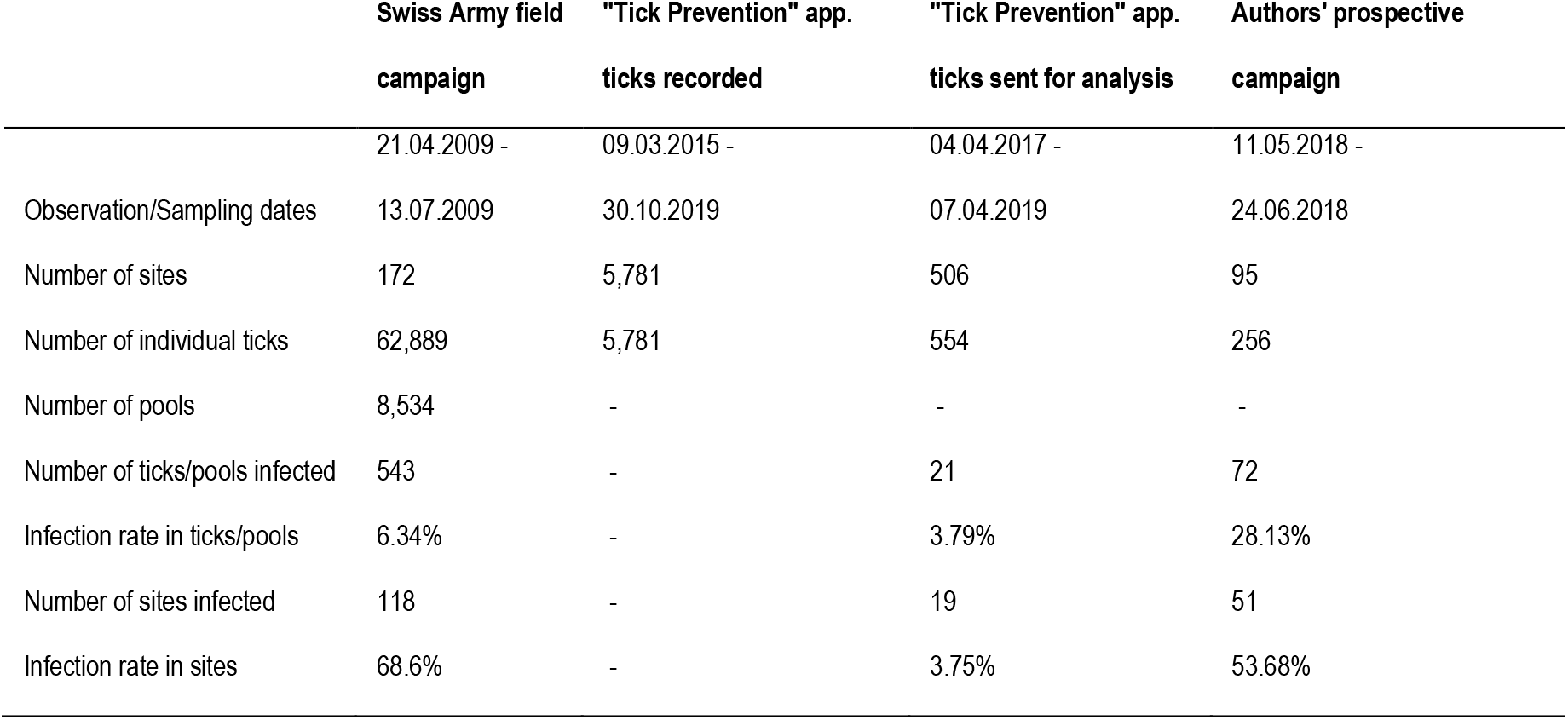
Characteristics of the three data sources regarding Ixodes ricinus occurrences and infection by Chlamydiales bacteria. The data obtained via the Tick Prevention app are divided into two datasets (column 2 and 3). The first dataset (column 2) corresponds to tick locations recorded on the app. including a majority of ticks for which no information regarding Chlamydiales bacteria were available. This dataset was used in the modelling of the distribution of Ixodes ricinus only. The second dataset (column 3, which represents a subset of dataset listed in column 2) contains some ticks that were sent to laboratory for the analysis of Chlamydiales. This dataset was therefore used in the modelling of Chlamydiales distribution. Data from the two other sources (column 1 and 4) were used both for the modelling of I. ricinus and Chlamydiales.

### Environmental data

To characterise the environmental conditions potentially influencing the spatial distribution of *Ixodes ricinus* and *Chlamydiales*, several information were retrieved for the whole Switzerland territory regarding 1) the morphometry 2) the land cover and 3) the climate.

To characterise the **morphometry** of each data point site, seven indicators were derived from the digital elevation model provided by the USGS/NASA SRTM data version 4.1, at a 90m-resolution (Jarvis *et al.*, 2008). The chosen indicators were computed using the SAGA GIS 2.3.2 software (Conrad *et al.*, 2015) and represent: slope, aspect, general curvature, morphometric protection index, terrain ruggedness, sky-view factor and topographic wetness. The definition of each of these indicators and the exact procedure followed to derive them are detailed in Supp. File 3.

To characterise the **land cover**, we first used the land cover statistics from the Swiss Federal Statistical Office (OFS, 2017). From this dataset we retrieved the classification of each Swiss hectare into six land cover types representative of the period 2004-2009: artificial areas, grass and herb vegetation, brush vegetation, tree vegetation, bare land and watery areas. To better classify forest type, we computed in R (R Development Core Team, 2008) the percentage of coniferous in each forest based on a dataset provided by the OFS at a 25-m resolution which classifies the forests of Switzerland in four classes : pure coniferous, mixed coniferous, mixed broadleaved and pure broadleaved (OFS, 2013). Secondly, we retrieved the vector landscape model swissTLM3D 2016 from the Swiss Federal Office of Topography (O’Sullivan *et al.*, 2008) and we use the function “Proximity” in the QGIS 2.14.7 software (QGIS Development Team, 2016) to derive four indices characterising the minimal Euclidean distance to watery areas: distance to wetland, to watercourses, to stagnant water and to any watery elements. Thirdly, we retrieved the 16-days composite Normalised Difference Vegetation Index (NDVI) available in the MODIS Satellite products at a 250m-resolution (Huete *et al.*, 1999), from which we derived in R the average, minimum, maximum and range of monthly mean NDVI. More details regarding all those land cover data and the derived indicators are also available in Supp. File. 3.

Finally, several indicators were computed to summarise the **climatic** conditions of each data point site. They were derived from monthly temperature (average, minimal and maximal) and sum of precipitation grids computed at a 100m-resolution by the Swiss Federal Institute for Forest, Snow and Landscape Research (www.wsl.ch), based on data from MeteoSwiss (www.meteoswiss.ch) and using the Daymet software (Thornton *et al.*, 1997). From these data, 31 indicators were derived to represent the climatic conditions during the period of interest and before sampling date (from 1 to 36 months preceding sampling date, see extraction chapter for more details). These indicators are presented in Supp. File 3 and they summarise 1) the values of the monthly mean, minimal and maximal temperature and sum of precipitation (8 indicators), 2) the variation of monthly temperature and precipitation (5 indicators), 3) the temperature of the warmest (resp. coldest) month (2 indicators) and 4) the temperature and precipitation of the three consecutive warmest (resp. coldest, wettest, driest) months (16 indicators). In addition, grids of the daily maximum and minimum temperature values at a 1km-resolution were obtained from MeteoSwiss. From these datasets, we estimated the daily saturated and ambient vapour pressure using the Tetens formula (Murray, 1966) and by approximating the temperature at dew point by the minimum temperature (Running *et al.*, 1987). We used them to compute the daily relative humidity and to derive 22 indicators summarising the monthly (9 indicators) and daily (13 indicators) values of relative humidity. All these climatic predictors were computed in R, with the detailed procedure presented in Supp. File 3. In total, this resulted in 77 environmental indicators, each of which were resampled to a final spatial resolution of 100 m.

### Data extraction

The values of the 77 environmental predictors were extracted for each data point site (tick occurrence) according to their coordinates using the function “extract” from the R “raster” package. The climatic and NDVI variables were retrieved as a function of the sampling dates. To assess the influence of the conditions before sampling, we retrieved these variables for 1 month, 3 months, 6 months, 1 year, 2 years and 3 years before sampling date. For the other stable predictors such as morphometric predictors, land cover type, percentage of coniferous in forest and distances to watery areas one single extraction was used for all sampling dates over the period of analysis (from 2009 to 2019).

To assess the influence of the environmental conditions surrounding the sampling points, for each environmental predictor we also computed the mean value observed in square buffers centred on the sampling point, with radius of 100 m, 200 m, 500 m, 700 m, 1 km and 1.5 km. Raster layers were also computed for each of these indicators, with every buffer radius and time period, for June months from 2009 to 2019. For each pixel, the computation of mean values considering a square buffer around the pixel was done with a moving-window procedure implemented in R, based on the “focal” function from the “raster” package.

Finally, we also extracted all predictors for a randomly generated data set (to test it against sampling data, see hereafter). This generated data set is composed by sites with 10’000 coordinates randomly localised in Switzerland, for which dates were selected randomly within the distribution of observed sampling dates (Supp. File 4).

### Ixodes ricinus modelling

#### Selection of environmental variables

The species distribution models were successively derived using the variables extracted for each combination of buffer radius (100 m, 200 m, 500 m, 700 m, 1 km and 1.5 km) and time period (1 month, 3 months, 6 months, 1 year, 2 years and 3 years). In addition, to select the most significant combination of buffer radius and time period individually for each variable, we performed a Student T-test to identify the variables that best discriminate the tick’s presences from random points. The computation was done using the function “t.test” in R and variables were considered as significant if the p-value of the T-test was lower than 0.01 after a Bonferroni correction for multiple comparisons. For each variable, we then kept only the combination of buffer radius and time period showing the highest T-value. A “combination” model was then derived using this “combination” set of variable.

As some environmental variables considered might be correlated, we used two methods to pre-select uncorrelated environmental predictors. In the first one, we run a Principal Component Analysis (PCA) on the variables to retrieved independent components. The coordinates of the PCA-components were then used as environmental predictors to run the species distribution model. In the second method, for each pair of variables showing a Pearson correlation higher than 0.8, we kept only the variable with the highest T-value in the T-test previously computed. In addition, we successively removed the variables inducing the highest inflation factor (VIF) computed with the R function “vif”, until the highest VIF value was lower than 10. Only the remaining variables were used to train the model.

#### Maxent Modelling

Species distribution modelling was performed using the Maxent algorithm (Phillips *et al.*, 2006) implemented in the R package “maxnet” (Phillips *et al.*, 2017). Maxent estimates a suitability index which is proportional to the probability of presence of the species knowing the environmental conditions of a site of interest (Elith *et al.*, 2010). The computation requires the values of environmental predictors observed on sites where presence was recorded and on background locations (i.e. locations representative of the entire study area). The model was trained with all *Ixodes ricinus* occurrences available for years 2009 to 2017 and the occurrences from the 2018 prospective campaign. This represents a total of 2293 presence points. The occurrences reported by the users of the Tick Prevention app. in 2018 and 2019 with 3751 presence points were kept as an independent dataset used to test the models.

Since the performance of the Maxent models is known to be influenced notably by the background point selection, environmental variable selection, features types and regularisation parameters (Lobo and Tognelli, 2011; Barbet-Massin *et al.*, 2012; Merow *et al.*, 2013; Hallgren *et al.*, 2019), we tested different alternatives regarding them. For the selection of background points, we tested two options: either we used the 10 000 points randomly selected in the Swiss territory or we used only the random points situated below 1500 m in altitude, where tick occurrence is more likely. For the environmental variables, we used the two procedures to derive uncorrelated set of variables, i.e. the coordinates of the PCA components and the variables filtered by the previously described method based on Pearson correlation and variance inflation factor. Moreover, when using the PCA components, we considered either all components of the PCA or only the components needed to retain 50% of the variance, resp. 70%, 80%, 90% or 95%. For the feature types, we tested the use of linear features only, or the combination of linear and product, linear and quadratic or linear, product and quadratic together. Finally, we varied the regularisation constant parameters with values equal to 1, 2, 5 or 10.

In order to perform a cross-validation procedure, we used 75% of the occurrences and background points to train the model and kept 25% to test it. The training and testing occurrences were selected randomly and 20 different runs were computed. All models were projected using the “cloglog” scaled output (Phillips, 2017), interpreted in terms of suitability index to avoid making assumptions regarding the prevalence of the species.

#### Model evaluation

The models were compared based on four criteria. First the Area under the Receiver Operating curve (AUC) (Fielding and Bell, 1997) was computed on the testing dataset. The mean value of AUC_test_ over the 20 runs was used as a measure of discrimination power. The AUC is a measure commonly used for the evaluation of species distribution models (Manel *et al.*, 2001; Elith *et al.*, 2006). It has the advantage to be threshold-independent, but needs to be used in combination with other evaluation parameters (Lobo *et al.*, 2008; Peterson *et al.*, 2008; Jiménez-Valverde, 2012). Therefore, we used as a second evaluation measure the omission error rate, which reflects the accuracy of the model. The computation of this rate requires the definition of a threshold value to classify the predictions into binary presences or absences. Based on the receiver operating curve, we chose the threshold which maximises the sum of specificity and sensitivity and therefore minimizes the misclassification rate (Kaivanto, 2008). Omission errors were computed both on the testing and independent (3751 points from 2018 and 2019) datasets. Finally, to avoid the selection of complex models, that would be difficult to interpret and probably prone to overfitting, we used a third evaluation measure that selected against models having high number of coefficients (following the principle of information criterion (Aho *et al.*, 2014))..

To combine the four evaluation parameters and select the most powerful model, we assigned four performance ranks to each model as a function of each evaluating parameter and we selected the model which minimises the sum of ranks. We then applied the best model to the raster layers to map the predicted suitability across entire Switzerland for June months from 2009 to 2019.

#### Identification of effective variables

In order to identify the environmental variables most contributing to the model, we implemented in R a jackknife procedure as proposed by Phillips (2017). For each environmental predictor, we computed the Maxent model with only this variable and calculated the corresponding AUC (AUC_only_). Variables leading to high values of AUC_only_ therefore contribute a lot to the model by themselves. Similarly, we successively computed models with all variables except the one under interest and we computed the corresponding AUC_without_. Predictors associated with high values of AUC_without_ were identified as containing important information that is not present in the other variables.

### Chlamydiales Modelling

#### Background dataset

To model the distribution of *Chlamydiales* bacteria within ticks, we used a similar procedure to that of *Ixodes ricinus*. The modelling was also done using Maxent, based on the 186 occurrence points available for 2009 and 2018. As for *I. ricinus*, the modelling required the definition of background data. Since we are interested by the probability to find *Chlamydiales* within ticks, background points have to represent the environmental conditions of the ecological niche for the tick. Consequently, we built a background dataset in two steps. First, we selected the points where ticks have been observed and analysed for the presence of *Chlamydiales*, but being negative (374 points). Secondly, in order to avoid a model discriminating presences from background due to differences in sampling dates, we completed the background dataset such to have a similar distribution of sampling months and sampling years as in the presence dataset (Supp. File 4). This was achieved by selecting random points within areas predicted to be suitable for ticks, based on the suitability predicted by the models previously derived for *Ixodes ricinus*. The final background dataset contains 1028 data points.

#### Variable selection and modelling

The same procedure was then applied as for the modelling of the tick’s suitability: 1) computation of a T-test to select a “combination” dataset of environmental variables, 2) selection of uncorrelated variables with either a PCA or a correlation/VIF procedure, 3) run of Maxent models by testing various parameters (method to select uncorrelated variables, feature types and regularisation parameters). In order to build models for the suitability of *Chlamydiales* within areas suitable for ticks, the predicted suitability for *Chlamydiales* obtained by the Maxent model was then multiplied by the suitability obtained for *I. ricinus*.

As for *I. ricinus*, twenty runs were computed for each model, using 75% of the data to train the model and 25% to test it. The ranking procedure used to evaluate the models was slightly different to the one used for the tick. The AUC_test_ and the number of coefficients were used similarly, but the omission rates on testing and independent datasets were replaced by two other indicators 1) the difference between the mean of suitability values predicted on occurrences sites in 2009 and the mean suitability predicted on sites without *Chlamydiales* in 2009 and 2) the same difference for 2018. Indeed, even if sites where no *Chlamydiales* were found could not be considered as proper absences, we suspected the probability to find *Chlamydiales* to be lower on these sites. A model showing a lower suitability in areas where *Chlamydiales* were not identified as compared to occurrence sites would therefore be considered as more performant.

## Results

### Ixodes ricinus modelling

#### Best model

Among the 56 models tested with various parameters, the best one, according to the ranking procedure, was obtained with the following parameters: 1) background points selected below 1500 m in altitude (corresponding to 6049/10 000 points), 2) a PCA procedure to avoid correlated variables, with the components selected to retained 95% of the variance, 3) a combination of linear and quadratic features and 4) a value of 5 for the regularisation constant parameter. Details of the models tested, and their corresponding evaluation parameters, are available in Supp. File 5. These parameters were then used to test the influence of the choice of buffer radius and time period on the performance of the models. Figure 1 shows the AUC_test_ and sum of ranks obtained for each combination. According to these results, the best model was obtained by extracting the environmental variables in a buffer with a 100-m radius around the sampling point and for the 2 years (24 months) preceding the sampling date. Note that the performance of the “combination” model was very close, as well as the performance of models obtained with an extraction for the 3 years preceding sampling date and a buffer radius of 100 m, or for the two years preceding sampling date with a 200 m buffer. Moreover, we observed for each buffer radius, that the models were more performant when considering the variables extracted for the 2 or 3 years previous sampling date, instead of considering the conditions of the current year or even shorter time period. Similarly, the models obtained by extracting the variables within buffers of 100 m or 200 m radius always outperformed the other models. Performance of models with variables extracted at the sampling coordinates only (radius = 0m) was much lower than any buffer model, even those with a radius larger than 500 m. We retained the best model with variables extracted in a 100 m-radius buffer and for the two years preceding the sampling date (Figure 1). The global AUC obtained (with both the training and testing data) is 0.794 and the mean AUC_test_ obtained through the 20 runs is of 0.789. The threshold maximising the sum of sensitivity and specificity equals 0.59. Using this threshold, the average omission error on the testing dataset reach 23% and the omission rate on the independent dataset is 11%. The model estimated 31 non-negative coefficients. The median predicted suitability on all occurrences used in the model is 0.74 and the median suitability on independent occurrences from 2018 and 2019 is 0.88.

**Figure 1:**
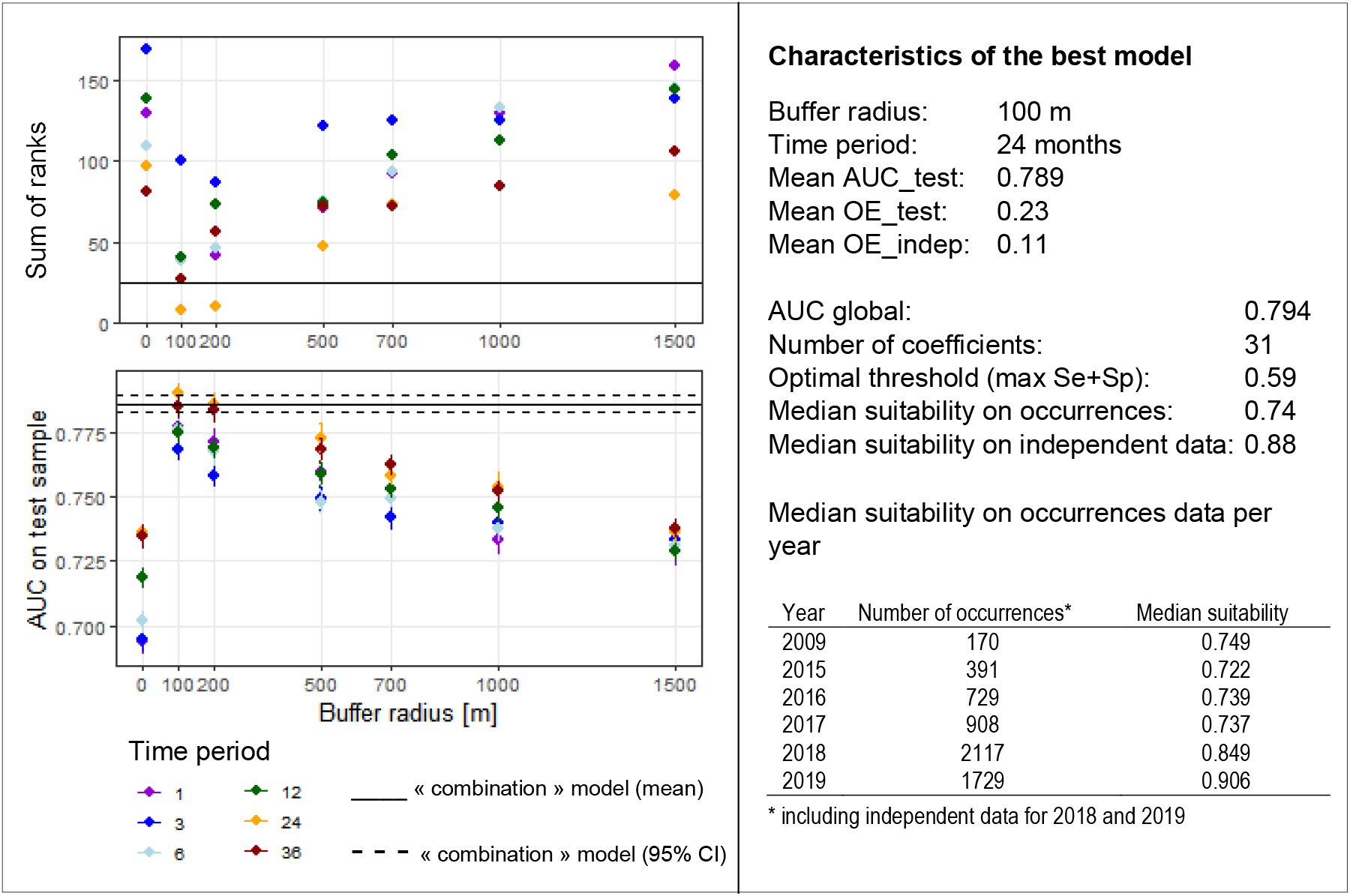
Performance of models predicting the suitability for *Ixodes ricinus*. (**Left**) Values of the AUC_test_ and the sum of ranks as a function of the buffer radius and the time period considered for the extraction of the environmental variables. For the AUC_test_, the points indicate the mean value computed through the 20 runs and the lines correspond to the 95% confidence intervals. (**Right**) Characteristics of the best model chosen according to best values on the graphics on the left. OE_test is the omission error on the test samples and OE_indep the omission errors on the independent additional data available for 2018 and 2019.

#### Effective variables

The four variables containing the largest amount of important information not available in the other variables (lowest AUC_without_) were: the dimension 1 (AUC_without_=0.748), dimension 12 (0.776), dimension 8 (0.780) and dimension 5 (0.784) (using jackknife procedure, Figure 2) indicated that the four variables containing the largest amount of important information by themselves (highest AUC_only_) were: the first dimension of the PCA (AUC_only_=0.641), the dimension 12 (0.617), dimension 21 (0.591) and dimension 8 (0.582).

**Figure 2:**
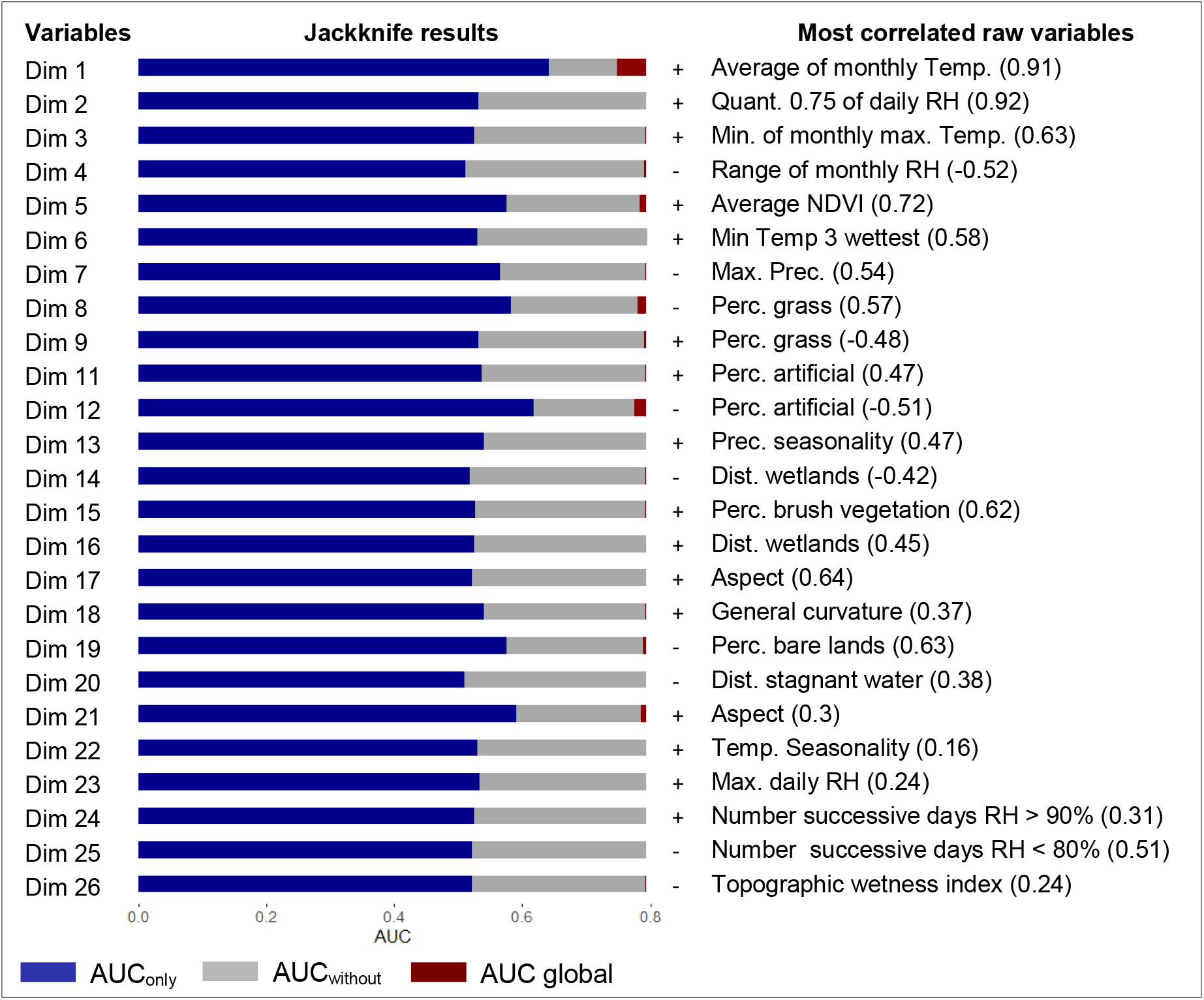
Jackknife results for the best model predicting the suitability of *Ixodes ricinus*. The variables Dim1 – Dim26 correspond to the components of the PCA needed to retain 95% of the variance. The column with +/− indicates the type of association between the component and the presence of *Ixodes ricinus* (with a positive association, the higher the value of the PCA dimension, the higher the suitability for ticks). The last column shows the raw environmental variable most correlated to the PCA dimension, with the value of the correlation indicated in parenthesis (Temp. = Temperature, RH = Relative Humidity, Quant. = Quantile, Prec. = Precipitation, Perc. = Percentage)

The dimension 1 of the PCA is strongly positively correlated with average of the monthly mean temperatures (r=0.91) and indicates that presence of *Ixodes ricinus* is favoured by higher mean temperature. Dimension 8 is moderately correlated with the percentage of herbs and grass vegetation (r=0.57) and the mean temperature during the three consecutive driest months (r=0.40). Its negative coefficient indicates that a higher percentage of herb and grass vegetation or higher temperature values during the driest months are less favourable for the presence of ticks. Dimension 12 is moderately negatively correlated with the percentage of artificial surfaces (r=−0.51) and positively correlated with the range of monthly NDVI (r=0.35). This dimension is also negatively associated with the suitability for ticks, indicating that a higher percentage of artificial surfaces and a lower range of NDVI values are more favourable for *I. ricinus* presence. Finally, the dimension 5 is positively correlated with the mean monthly NDVI (r=0.72), the minimum and maximum NDVI (r=0.55 and 0.52) and is negatively correlated with the percentage of watery areas (r=−0.56). Its positive coefficient indicates that the areas with higher NDVI values and less water are more favourable for ticks.

#### Distribution maps

The maps of the distribution of *Ixodes ricinus* with values of suitability index predicted by the model across Switzerland for June 2009 and June 2018 are shown on Figure 3. The corresponding projections for June 2015, 2016, 2017 and 2019 are available in Supp. File 6. Results for June 2009 shows that 16% of the Swiss territory is predicted suitable for the presence of *Ixodes ricinus*, when using the threshold maximising the sum of specificity and sensitivity (threshold = 0.59). The suitable areas are mainly localized in land covered by tree vegetation (48.6 % of all suitable areas), however 26.6% are observed on hectares statistically classified as artificial surfaces. In addition, most of suitable area lied between 500 and 1000 m in altitude (53.04%) or below 500 m (46.5%). Only 8.4 % of the favourable area is found above 1000 m in altitude.

**Figure 3:**
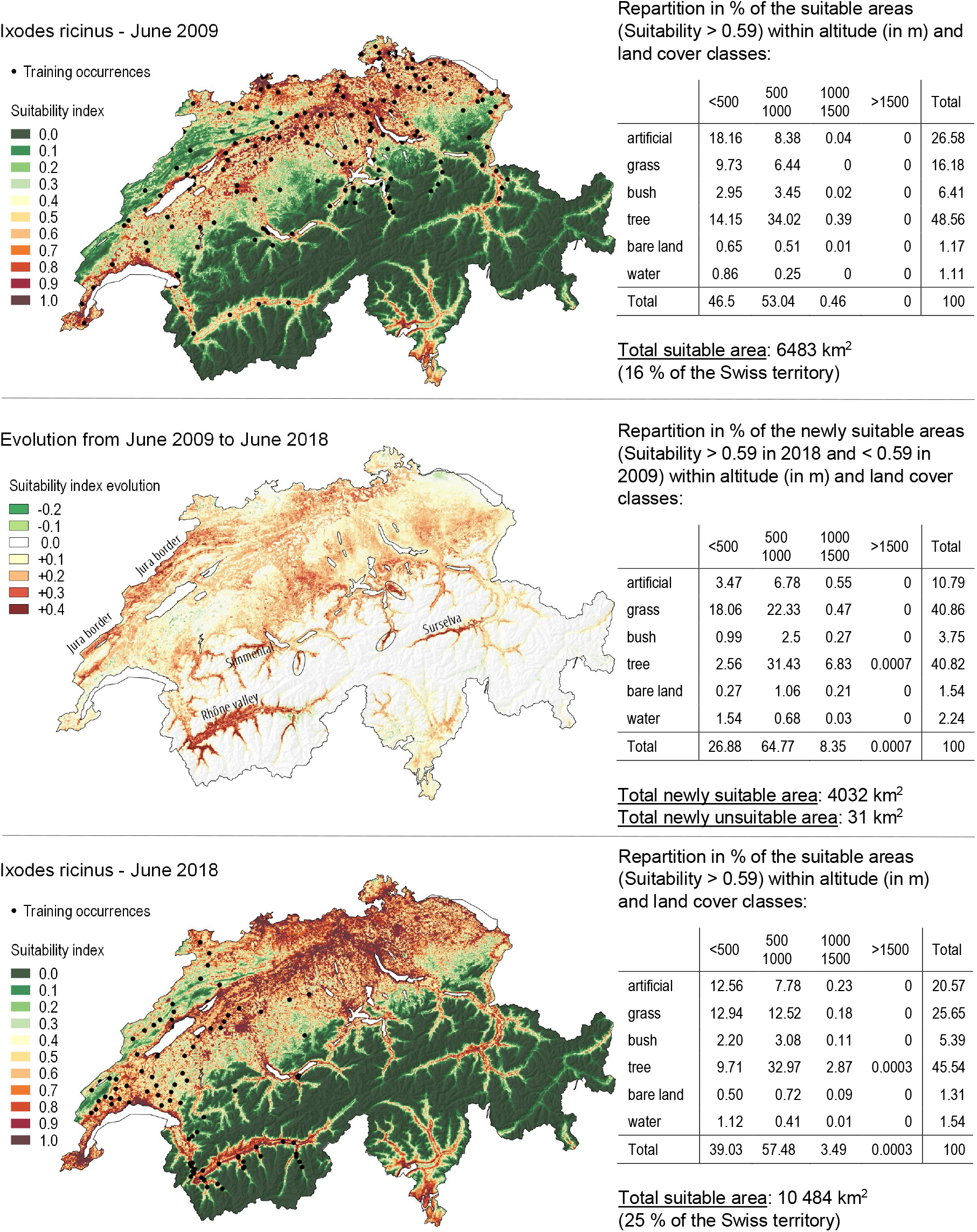
Suitability maps for *Ixodes ricinus*. Suitability map for *Ixodes ricinus* in June 2009 (upper panel) and June 2018 (lower panel) as predicted by the best model (i.e. with environmental variables extracted with a 100m-radius buffer and for the two years preceding sampling date). The area concerned by the transition in suitability are represented in the intermediate panel.

In June 2018, 25% of the Swiss territory is predicted suitable for *Ixodes ricinus* (considering the threshold of 0.59). Between June 2009 and 2018, the predicted suitable area increased by more than 4000 km^2^ as shown in Figure 3 and only 31 km^2^ became unsuitable. The increased suitability is particularly pronounced in the Rhône Valley (Valais), in Surselva, in Simmental, in the Jura border and in other lateral valleys of medium to high altitude (circles on the map). The evolution of the PCA components from 2009 to 2018 in these areas shows that the increase in suitability is generally associated with an increase of the values of Dimension 1 (warmer temperature), an increase of Dimension 5 (higher NDVI values), a decrease of Dimension 12 (lower range of NDVI values), and a decrease of Dimension 8 (temperature during driest months) in Valais and Jura (whereas this last dimension shows an increase of the values in Grisons). The new suitable areas concerned mainly grass and tree vegetation (40.8% each) with a large proportion (64.8%) located at an altitude between 500 and 1000 m (corresponding for example to the altitude of the suited hectares in Jura border or Rhône valley). An increase of suitable areas mainly in forests was also observed between 1000 and 1500 m (8%).The model also predicted suitable areas above 1500 m. These results therefore highlighted a spread of the favourable areas towards higher altitude.

The distribution maps of *Ixodes ricinus* for the years 2015 to 2017 (Supp. File 6) indicate a constant and drastic increase in suitability which is highest between 2017 and 2018. Indeed, 15.7% of the Swiss territory was predicted as suitable in 2009, 16.8% in 2015, 16.2% in 2016, 17.6% in 2017 and 25.4% in 2018 (by considering the threshold of 0.59 for suitable areas). Moreover, the map computed for 2019 predicted important increase from 2018 to 2019, with 35% of the Swiss territory being predicted as suitable in 2019. The spread towards higher altitude was also observed between 2018 and 2019, with a maximal altitude for the favourable areas that reached 1595 m in 2019. The results indicate that since 2018, there is a relatively high probability that ticks reach such altitudes.

### Chlamydiales modelling

#### Best model

The best model for *Chlamydiales* bacteria, among the 60 models tested with various parameters, was obtained with the following parameters: 1) the “correlation-VIF” procedure to select uncorrelated variables, 2) a combination of linear and quadratic features and 3) a value of 1 for the regularisation constant parameter. The details of all models tested and their corresponding evaluation parameters are available in Supp. File 7. As for the modelling of *Ixodes ricinus*, we then tested the influence of the choice of buffer radius and time period on the performance of the models. Figure 4 shows the AUC_test_ and sum of ranks obtained for each combination. According to these results, the “combination” model outperformed the other models. Unlike the results obtained for *Ixodes ricinus* the models for *Chlamydiales* performed better when the variables are extracted for the three- or six-months preceding sampling date than when considering two or three years before sampling (Figure 4). In addition, the influence of buffer radius seems to be much less pronounced than for the tick models. Accordingly, we retained the “combination” model. This model used 17 uncorrelated variables selected based on the “correlation/VIF” procedure. The list of these variables, as well as the results of the T-test are available in Supp. File 8. As the “combination” model aims to retain for each variable the best combination of buffer radius and time period, not all variables are selected using the same buffer radius or time period. Interestingly, we observed that the variables used in the model involved either buffer radius smaller or equal to 200 m, or superior to 1 km (Supp. File 8). The characteristics of the model are summarised on the right of Figure 4. The global AUC (with both training and testing occurrences) is 0.78 and the mean AUC_test_ obtained through the 20 runs is of 0.74. The threshold maximising the sum of sensitivity and specificity equals 0.3. The mean suitability for *Chlamydiales* occurrence in 2009 is 0.47 and the mean suitability for sites where *Chlamydiales* where not identified in 2009 is 0.37. For 2018, the mean suitability on presence points is 0.46 and the suitability on sites where no *Chlamydiales* were identified is 0.15. The model estimated 35 non-negative coefficients.

**Figure 4:**
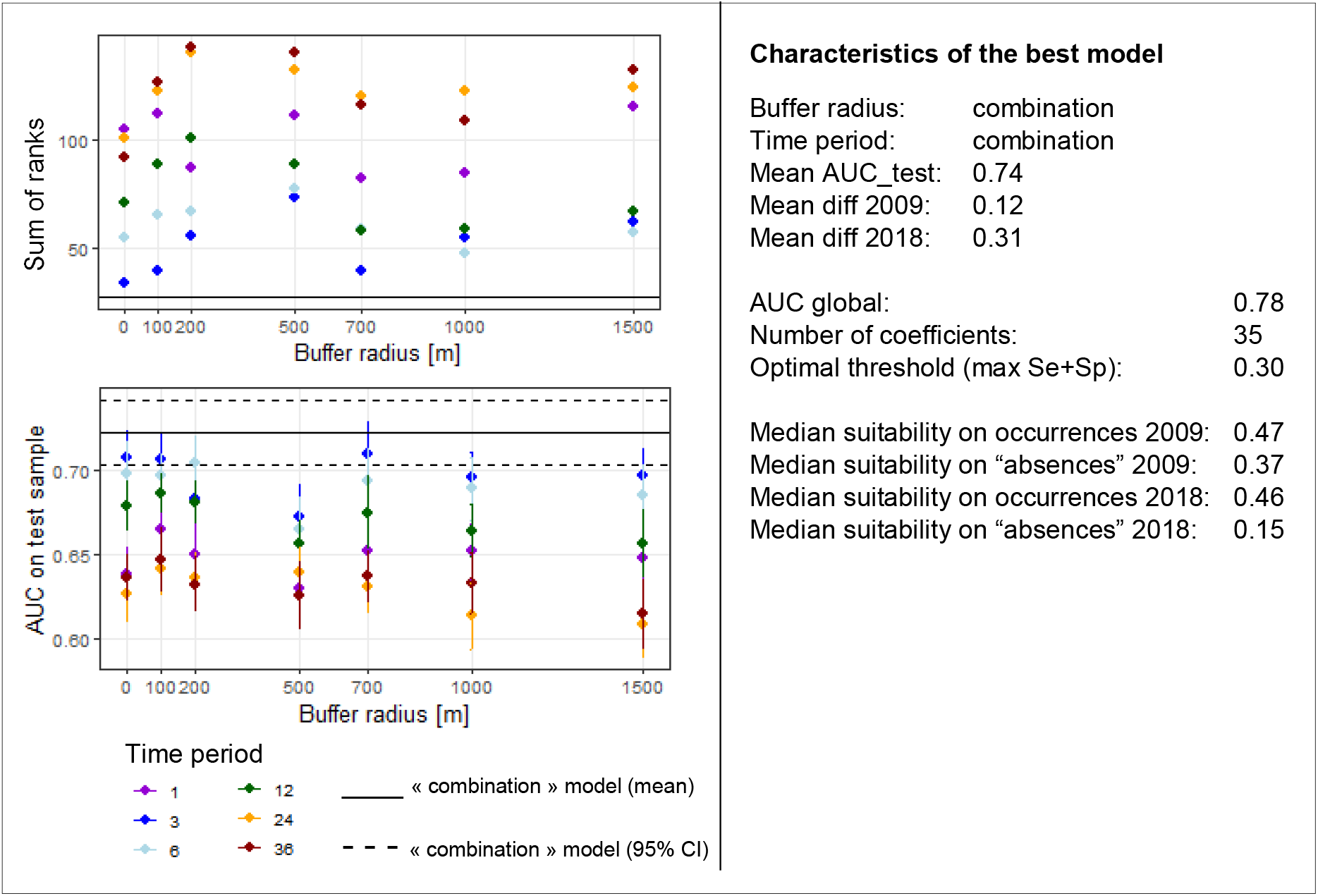
Performance of models predicting the suitability for *Chlamydiales*. (Left) Values of the AUC_test_ and the sum of ranks as a function of the buffer radius and the time period considered for the extraction of the environmental variables. For the AUC_test_, the points indicate the mean value computed over the 20 runs and the lines correspond to the 95% confidence intervals. (Right) Characteristics of the best model chosen according to the graphics on the left. Mean diff 2009 (resp. 2018) is the average difference between the mean suitability values predicted on *Chlamydiales* occurrences points and on locations where no *Chlamydiales* were identified in 2009 (resp. 2018).

#### Effective variables

The four variables containing the highest amount of important information that are not available in the other variables (lowest AUC_without_) are (Figure 5): the percentage of tree vegetation in a 100 m buffer (AUC_without_ = 0.75), the coordinates (no buffer) number of successive days with a relative humidity inferior to 80% during the 3 months preceding sampling (0.77) or inferior to 70% during the 6 months preceding sampling (0.77) and the distance to wetlands within a buffer of 1km (0.77). The four variables containing the highest amount of important information by themselves (highest AUC_only_) are: the percentage of artificial surfaces in a 100 m buffer (AUC_only_ = 0.59), the number of days with a relative humidity superior to 90% in a 200 m buffer during the two years preceding sampling date (0.57), the precipitation of the three coldest months in a 1.5 km buffer during the two years preceding sampling (0.55) and the percentage of tree vegetation in a 100 m buffer around the sampling point (0.55).

**Figure 5:**
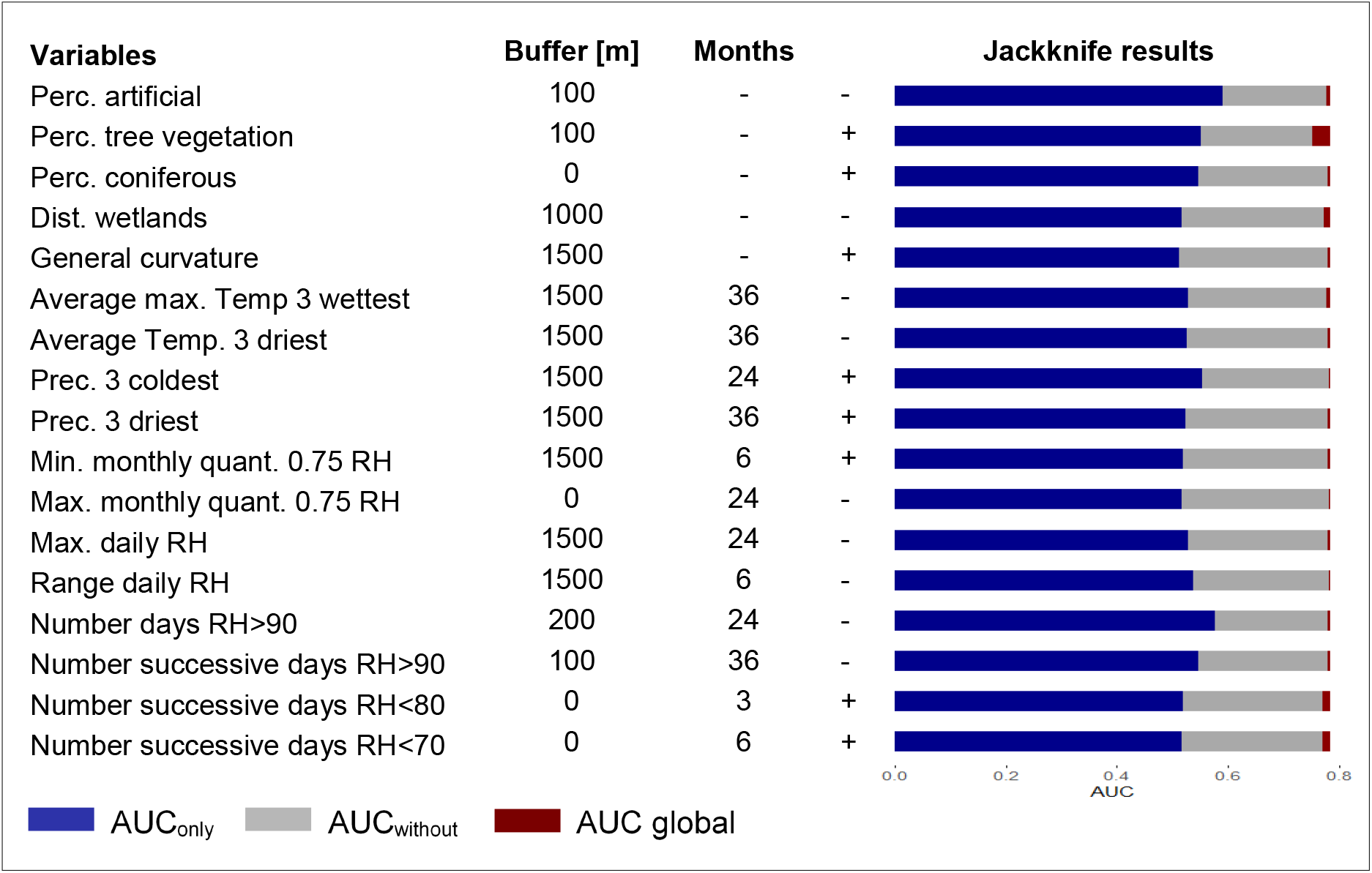
Jackknife results for the best model predicting the suitability of *Chlamydiales*. The column “Buffer” indicates the buffer radius around the sampling point and “Months” the number of months before sampling date. The column with +/− indicates the type of association between the variable and the presence of *Chlamydiales* (with a positive association, the higher the value of the variable, the higher the suitability for *Chlamydiales*). Perc. = Percentage, Temp. = Temperature, Prec. = Precipitation, quant. 0.75 = quantile 0.75, RH = Relative Humidity.

The conditions favourable for *Chlamydiales* are thus characterised by: a lower percentage of artificial surfaces around the sampling point (7.8% in average for the occurrences locations in a 100m-buffer versus 16.8% for the background locations), a higher percentage of tree vegetation (62.8% versus 53.1%), a lower number of days with a relative humidity superior to 90% during the two years preceding sampling date (21.1 versus 25.2), a highest amount of precipitation during the coldest months (24.15mm versus 20.7mm), a higher number of successive days with a relative humidity inferior to 80% during the three previous months (29.7 versus 27.1) and lower than 70% during the 6 previous months (16 versus 14.4) and finally a shorter distance to wetlands (2.5 km versus 3.1km).

#### Distribution maps

The distribution maps of *Chlamydiales* with values of suitability predicted by the model across Switzerland for June 2009 and June 2018 are shown on Figure 6. In June 2009, 8% of the Swiss territory is predicted as favourable for *Chlamydiales* bacteria (using the threshold maximising the sum of sensitivity and specificity). As the niche of the bacteria is nested within the niche of the tick, modelling *Chlamydiales* bacteria suitability involved a multiplication by the suitability results for *Ixodes ricinus*. Therefore, the areas predicted to be unfavourable for the presence of the tick species are also predicted as weakly suitable for *Chlamydiales.* On the contrary, some areas predicted to be highly favourable for the presence of *Ixodes ricinus* on Figure 3 did not match and showed very low values on Figure 6. This is the case for the areas situated within urban settlements, in which a large portion was predicted to be suitable for ticks but not for *Chlamydiales*. Indeed, the distribution of the favourable areas within the various categories of land cover classes indicates that they are essentially observed in natural areas, covered either by tree (74%) or grass (12%) vegetation, and only 4% of them are observed in regions characterised by a large portion of artificial elements. When considering the altitudinal distribution, areas favourable for *Chlamydiales* seem to be essentially predicted in forest suitable for ticks, between 500 and 1000 m in altitude. However, due to other factors influencing the model, notably the climatic conditions, 52% of those forests are also predicted to be unfavourable for the bacteria.

**Figure 6:**
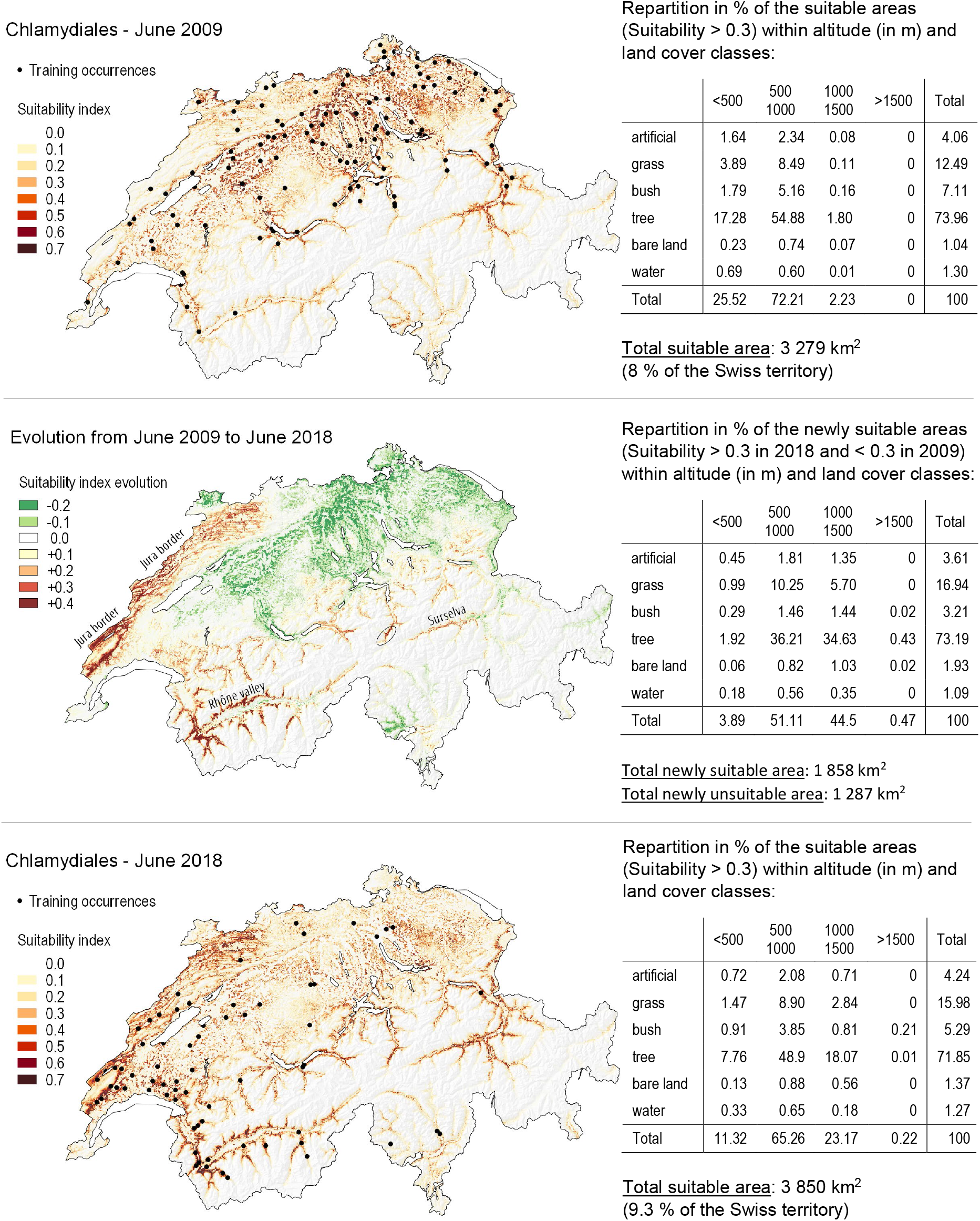
Suitability maps for *Chlamydiales*. Suitability map for *Chlamydiales* in June 2009 (upper panel) and June 2018 (lower panel) as predicted by the best model (i.e. with “composition” set of environmental variables). The area concerned by the transition in suitability are represented in the intermediate panel.

In June 2018, 9% of the Swiss territory is predicted as suitable for the presence of *Chlamydiales*. Between June 2009 and 2018, more than 1850 km^2^ are newly suitable for *Chlamydiales* as shown in Figure 6. Some regions showing a sharp increase in suitability values (more than 0.4). However, more than 1,300 km^2^ is also becoming unsuitable. In 2018, the proportion of suitable area within land cover classes is close to what observed in 2009, with however a clear spread towards higher altitude, with 23% of the favourable areas localised between 1000 and 1500 m, versus 2% only in 2009. Newly suitable area match those of *Ixodes ricinus* on Figure 3 (Rhône valley, Surselva, Jura border). The spread of favourable areas towards higher altitude is also predicted, with 45% of the newly suitable hectares being localised between 1000 and 1500 m. Loss of suitable area mainly occurred in the North-West part of Switzerland and appear to be associated with a decrease in precipitation during the three coldest months and a decrease of the successive number of days with a relative humidity inferior to 70% during the 6 previous months (15th of December 2017 to 15^th^ of June 2018 as compared to 15^th^ of December 2008 to 15^th^ of June 2009).

## Discussion

### Expansion of Ixodes ricinus and Chlamydiales in Switzerland

Distribution maps for ticks and bacteria from 2009 to 2019 highlighted an extension of the suitable areas for both species and a spread towards higher altitude. *Ixodes ricinus* expended from 16% to 25% of the Swiss territory, and a subsequent extension for *Chlamydiales* bacteria is observed from 8% to 9.3%. *Ixodes ricinus* expansion occurred all over the Swiss Plateau and toward higher altitude in the alpine valleys and was more extended in the South-West. Newly available habitat concerned mostly grass and forest areas. Extension of *Chlamydiales* followed similar trends, restricted to forest areas. As *Ixodes ricinus* presence is favoured by higher temperature, we might expect that, in the future, this expansion might continue following global warming with some limitation by dryer conditions at lower altitude

Our results agrees with the observed increased cases of tick-borne encephalitis (TBE) in Switzerland, that spread from eastern to western part of Switzerland (de Vallière and Cometta, 2006), leading to the extension of the vaccination recommendation (OFSP, 2013, 2019). Similar tick’s expansions towards higher altitudes were observed in other European countries during the last decades (Daniel *et al.*, 2003; Skarphédinsson *et al.*, 2005; Jore *et al.*, 2011), notably in association with milder winters and extended spring and autumn seasons (Lindgren E *et al.*, 2000; Medlock *et al.*, 2013).

### Variables explaining I. ricinus distribution

The effective variables identified by our model are related to temperature and humidity, which reflects well the tick’s ecology. We found that a high temperature favours *Ixodes ricinus*, in agreement with previous studies (Estrada-peña, 1999; Porretta *et al.*, 2013). However, our analysis indicated that this relationship does not hold during driest months. This can be explained by an increased evaporation of the soil humidity under warmer temperature, thus accentuating the desiccation risk for ticks (McCoy and Boulanger, 2015). The NDVI variables, an important contribution to our model, are indicators of physiological plant activity and have often been shown to be powerful for modelling the presence of ticks as they reflect humidity conditions (Estrada-peña, 1999; McCoy and Boulanger, 2015). Nevertheless, our results indicated that the ambient relative humidity variables showed limited effect on the model. They may thus constitute a less precise predictor of soil humidity than the combination of NDVI variables with temperature and land cover indicators. Surprisingly, our results also showed that *I. ricinus* presence is favoured by a higher percentage of artificial surfaces. This might relate to an overrepresentation of ticks collected in vegetated areas situated within urban settlements or close to roads. Indeed, we expect a sampling bias as many tick occurrences comes from the Tick Prevention App., in which users provide tick locations that are likely biased towards areas closer to roads or paths and thus artificial surfaces. Moreover, the other tick occurrences, either provided by the army field campaign in 2009 or by the prospective campaign in 2018, were collected essentially in forests or close to their borders. On the contrary, grass areas, often corresponding to agricultural fields, were not sampled by the two field campaigns and were also probably less explored by the users of the application, since people are less likely to visit these areas. This might explain why our model associated a low percentage of grass vegetation as favourable for *I. ricinus* and we might have an underestimation of the suitability index in some grass areas. Nevertheless, the presence of ticks in urban and suburban areas of Switzerland has already been reported (Rizzoli *et al.*, 2014; Oechslin *et al.*, 2017) and the presence of vegetated areas in urban settlement, or close to artificial surfaces (roads, paths, recreational areas) may constitute favourable habitats. In addition, even if we may expect some grass zones, especially at the forest border, to be highly favourable for ticks, in general, land pasture, open land and cultivated areas have been reported to be much less favourable than woodlands (Aeschlimann *et al.*, 1979; Huss and Braun-Fahrländer, 2007; McCoy and Boulanger, 2015). Finally, in agreement with previous studies (Estrada-Peña *et al.*, 2015; Hauser *et al.*, 2018), we observed that the morphometric parameters and the precipitation variables show little effect on the suitability for ticks.

### Variables for Chlamydiales spatial distribution

Identified effective variables for the presence of *Chlamydiales* may provide novel insights to the bacteria’s ecology. First, our results indicated that *Chlamydiales* are more likely present in ticks collected in forests or grass fields than in ticks collected close to artificial areas. The highest prevalence of *Chlamydiales* within natural areas could be explained by the presence of different hosts (likely rodents) on which ticks feed, with potentially a highest number of reservoir-competent hosts for *Chlamydiales* in natural areas. This may also relate to a higher tick abundance in natural areas, which is known to be associated with a higher prevalence of other pathogens in ticks (Aivelo *et al.*, 2019) but not for all tick pathogens (Oechslin *et al.*, 2017). Our results also showed that the presence of *Chlamydiales* bacteria is favoured by driest conditions (negatively associated with the number of days with a relative humidity superior to 90% and positively associated with the number of days with relative humidity inferior to 70%). High amount of precipitation during the coldest months also appeared to be favourable for the presence of *Chlamydiales*. Several suitable areas for *Chlamydiales* are predicted at an altitude higher than 1000 m, thus highest precipitation during the coldest months could be associated with largest snow amounts, preserving the soil from frost and leading to a highest tick’s survival (Lindgren *et al.*, 2006). Finally, a shorter distance to wetlands was also highlighted as a factor favouring the bacteria’s presence. Several *Chlamydiales* have been considered symbionts of amoebae (Kebbi-Beghdadi and Greub, 2014), which are free-living organisms usually found at the interface between water and soil, air or plants (Kebbi-Beghdadi and Greub, 2014). It is therefore likely that amoebae can be found in wetlands, which might favour the transmission of *Chlamydiales* to various animal hosts on which ticks feed.

*Chlamydiales* prevalence values were heterogeneous among our datasets. In 2009, ticks were collected in forests only and *Chlamydiales* were present in 68.6% of the sites visited with a low prevalence within pools (6.4%). Low prevalence was also observed in the ticks received by the users of the Tick Prevention App in 2018 and 2019 (3.79%). In 2018, the ticks sampled during the prospective campaign were also mainly collected in forest areas and *Chlamydiales* were present in 53.7% of the site but with much higher prevalence reaching 28.13%. This rate reflects values obtained in 2010 in one specific site in the Swiss Alps (Rarogne), where *Chlamydiales* prevalence rate of 28.1% was found in 192 pools collected in forests and meadows (Croxatto *et al.*, 2014). Differences between year 2009 and 2018 could be explained by a difference in the time and sampling areas (we excluded potential PCR contaminations, see Supp. File 9). As infected ticks were already present in most forest sites in 2009, spread of infection might have occurred between 2009 and 2018. Then, ticks from Tick Prevention App were collected in sites more closely related to artificial areas, which we have shown reduces the prevalence of the bacteria.

### On the importance of considering the spatial and temporal scale of the environmental variables

For *I. ricinus*, the most performant models are obtained when extracting the environmental variables in a buffer with a radius of 100 or 200 m (corresponding to an area of 9 ha to 25 ha around the sampling point). This can be explained by the ecology of the species. First, the establishment of a population of ticks will probably need a suitable area that is large enough. Moreover, the presence of ticks strongly depends on the presence of hosts, which disperse across larger areas and may thus be influenced by the climatic conditions observed at some distance. Our results also indicate that buffer radius larger than 500 m (corresponding to areas larger than 121 ha) are not improving our model. This might relate to the dispersal range of tick hosts, likely rodents, which is usually smaller (among the long dispersal hosts, the roe deer dispersal is estimated to cover around 50 and 100 hectares (Cederlund and Liberg, 1995)). In addition, the most performant models are obtained when considering the climatic conditions of the two- or three-years preceding sampling date. This time period appears to be relevant as it corresponds to the estimated duration of the life cycle of ticks (McCoy and Boulanger, 2015).

For the modelling of *Chlamydiales* bacteria, small buffer (≤200m) and a short time period (one year or less) is favourable for some variables, whereas for some others, to consider a larger buffer (1 km or 1.5 km) and a longer time period (2-3 years) is better. Some variables might be influencing locally the establishment of the tick species and the ability for the bacteria to colonize and/or reproduce within it, whereas other variables may be related to the interaction of the tick with the hosts on which it feeds, that may disperse in a larger area and thus be influenced by climatic conditions at a larger scale.

Our results thus highlighted the importance of considering the environment around the sampling point for a good variables estimation in species distribution model, while single point is commonly considered (Elith *et al.*, 2006; Williams *et al.*, 2015; Raghavan *et al.*, 2016, 2019, 2020; Sage *et al.*, 2017; Minigan *et al.*, 2018; Soucy *et al.*, 2018; Eisen *et al.*, 2018; Hadgu *et al.*, 2019). Our results also showed that the time period considered before the sampling date, with sliding windows, has a significant impact on the performance of the resulting models. This should be favour over using an average of the climatic conditions over the sampling period (Bradley *et al.*, 2010; Williams *et al.*, 2015) or any larger period of time (as Worldclim climatic data from 1950 to 2000 which are commonly used for species distribution modelling (Porfirio *et al.*, 2014; Manzoor *et al.*, 2018)). Previous studies already suggested the use of multi-grain approaches involving various spatial resolutions to consider variables affecting the presence of a species at different scales (Meyer and Thuiller, 2006; Meyer, 2007; Mertes *et al.*, 2020). This adds to the recommendation of using data based on species ecology rather than on availability (Mayer and Cameron, 2003; Meyer, 2007). In addition, our results showed that the temporal scale of the environmental predictors should be accounted for.

### Model performance

*Ixodes ricinus* distribution models are robust as they allowed a good discrimination between presences and randomly generated points and correctly predicted the presences of *I. ricinus* observed in an independent dataset. *Chlamydiales* distribution models are more difficult to validate due to the limited amount of data and poor knowledge regarding their distribution. Nevertheless, our model performed relatively well for the data collected in 2018 as most of the occurrence locations had higher suitability index than the locations where no *Chlamydiales* were identified. Year 2009 did not show such trend as many locations where no *Chlamydiales* were found were predicted as potentially suitable. This might be due to an absence of *Chlamydiales* colonisation of these sites at the sampling time despite favourable conditions.

Our investigations considered mainly environmental factors. However, other factors such as species interaction and species life history traits might influence the presence of both the ticks and their bacterial pathogens (Guisan and Zimmermann, 2000; Clay *et al.*, 2008; Estrada-Peña, 2008; Büchi and Vuilleumier, 2014; McCoy and Boulanger, 2015; Ehrmann *et al.*, 2018). Also, additional abiotic factors might play an important role, such as landscape fragmentation and barriers that can limit dispersal of ticks hosts (Estrada-Peña, 2008; McCoy and Boulanger, 2015) or disturbances that can drive local populations to extinction (Vuilleumier *et al.*, 2007).

The precision of our predictions is limited by the precision of the data used. The interpolated climatic grids used were produced based on weather stations measurements and thus contain interpolation uncertainties that may influence the models results (Guisan and Zimmermann, 2000). Also, with interpolated grids, the inherent collinearity and autocorrelation may lower the reliability of the results (Estrada-Peña *et al.*, 2015). Finally, the occurrences data are probably prone to sampling bias and do not represent a random sample of the population being studied, which can also influence the predictions (Araújo and Guisan, 2006; Merow *et al.*, 2013), probably leading to an overestimation of suitability index in urban and artificial areas as compared to natural ones.

## Conclusion

Both *Ixodes ricinus* and *Chlamydiales* are causing a potential threat to human health and their prevalence are currently increasing in Switzerland, with a strong expansion of ticks in forests but also in urban and suburban areas. Ticks’ expansion has already recently alarmed the Public Health Services (OFSP, 2019), and this expansion is predicted to continue in the future due to global warming. In this context, our results offer a unique tool to identify precisely locations where diseases are likely to spread, to colonize new sites and to increase in prevalence. Maps as developed here, and associated methods, could thus bring critical information for decision-makers to control tick-borne diseases and target prevention campaigns.

Our methodological framework allowed a coherent identification of environmental factors influencing the presence and distribution of both *Ixodes ricinus* tick and their *Chlamydiales* bacteria in Switzerland, and enabled the mapping of suitability evolution across Switzerland from 2009 to 2019. Our results highlighted an important increase of suitable areas for both species and predicted their extension towards higher altitude. Our investigations consist in the first exploratory analysis of the environmental factors influencing the presence of *Chlamydiales* bacteria within ticks in Switzerland, showing an application of species distribution models to study the nested niche of a parasite within the ecological niche of its host. Finally, our study demonstrated the importance of considering the spatial and temporal scale of the environmental variables used for species distribution models.

Spread of pathogens through a vector is at the origin of major epidemics and infectious diseases, and affects humans, wildlife, and agriculture. We proposed a methodological framework based on geographical system able to provide deep insights on factors affecting patterns of disease emergence by providing a better characterisation of the spatial distribution of their vectors. This method can be applied to a wide range of host-pathogen association to identify their spread and distribution, which is expected to bring critical information for a better understanding and control of pathogens.

## Acknowledgments

We thank Dr. Dirk Shmartz from the Swiss Federal Institute for Forest, Snow and Landscape Research, for computing and providing on demand the high resolution climate grids; Werner Tischhauser, Prof. Jürg Grunder and A&K Strategy for providing an access to the data of their smartphone application (Tick Prevention, https://zecke-tique-tick.ch); Rahel Ackermann-Gäumann for the tick data from the Swiss Army field campaign and Ludovic Pilloux for advices regarding the *Chlamydiales* dataset from this same field campaign.

## Code availability

The main R codes developed for this study are available on GitHub: https://github.com/estellerochat/SDM-Chlamydiales

